# Transient gamma events delineate somatosensory modality in S1

**DOI:** 10.1101/2023.03.30.534945

**Authors:** Christopher J. Black, Carl Y. Saab, David A. Borton

## Abstract

Gamma band activity localized to the primary somatosensory cortex (S1) in humans and animals is implicated in the higher order neural processing of painful and tactile stimuli. However, it is unclear if gamma band activity differs between these distinct somatosensory modalities. Here, we coupled a novel behavioral approach with chronic extracellular electrophysiology to investigate differences in S1 gamma band activity elicited by noxious and innocuous hind paw stimulation in transgenic mice. Like prior studies, we found that trial-averaged gamma power in S1 increased following both noxious and innocuous stimuli. However, on individual trials, we noticed that evoked gamma band activity was not a continuous oscillatory signal but a series of transient spectral events. Upon further analysis we found that there was a significantly higher incidence of these gamma band events following noxious stimulation than innocuous stimulation. These findings suggest that somatosensory stimuli may be represented by specific features of gamma band activity at the single trial level, which may provide insight to mechanisms underlying acute pain.

## 1. Introduction

Primary somatosensory cortex (S1) plays a significant role in pain perception. Historically, S1 has been attributed with representing the sensory-discriminative (intensity, location) component of pain, but S1 also helps regulate nocifensive reflexive behaviors through corticospinal projections [13], and modulates activity in cortical areas implicated in emotional pain processing through corticocortical projections [28,39]. This role in higher-order orchestration of pain processing makes S1 a prime target for understanding pain mechanisms in both acute and chronic pain states. Neural activity in the gamma band (30-100Hz) localized to S1 correlates with various dimensions of pain across species: Human electroencephalography (EEG) recordings over S1 have shown that gamma band oscillation (GBO) amplitude positively correlates with pain intensity [7,8,41], while electrocorticography (ECoG) recordings in rodents have shown that hyperalgesia positively correlates with GBO amplitude [38]. Furthermore, optogenetic induction of GBO in mouse S1 enhances nocifensive behavior [30]. These findings suggest that S1 oscillations in the gamma range relay noxious information to, and potentially coordinates neural activity in, higher-order brain circuits.

Although S1 gamma band activity is posited as a neural correlate of pain, similar signatures of activity in S1 are present during nonpainful touch. Innocuous somatosensory stimulation elicits increases in gamma power in both intracortical recordings in non-human primates [29] and EEG recordings in humans [17,18]. At the cellular level gamma activity generated by fast-spiking interneurons in sensory cortices enhances detection of tactile stimuli [25,27]. Together, these findings indicate a more generalized utility of gamma band activity in increasing saliency across somatosensory modalities as opposed to a specific neural correlate of pain.

An alternative interpretation of prior findings is that noxious and innocuous stimulations may differentially drive S1 circuitry to produce different types of gamma activity [15]. While research in humans using EEG has investigated the differences in S1 between noxious and innocuous responses at high frequency ranges [12,15], investigating differences between somatosensory modalities is confounded by the use of noxious stimuli that co-activate low-threshold sensory afferents carrying information about innocuous stimuli [10]. Preclinical rodent behaviors offer a solution to this issue with specific optogenetic strains that offer the ability to selectively activate subsets of A-delta and C-fiber afferents. These transgenic methods have also increased the reliability of preclinical pain measures by leveraging high-speed video and machine learning approaches to quantitatively characterize nocifensive reflexes within milliseconds from stimulus onset [1,19]. However, no study to date has examined differences in gamma activity between somatosensory modality at the intracortical level.

Therefore, we coupled peripheral optogenetics with quantitative behavioral measures of self-report to determine if a) nociceptive *or* tactile-specific stimulation evokes measurable differences in gamma band activity, and b) those modality-specific differences represent differential coding of sensory input. We developed an approach to investigate temporally precise and modality-specific sensory stimulation in transgenic mice, while implementing chronic electrophysiological recordings in S1. We identified features of transient gamma band activity that reflected distinct stimulus modalities. These results may prove useful in understanding the neural correlates of pain in the brain, and the potential to identify novel biomarkers of pain in clinical settings.

## 2. Methods

### 2.1 Transgenic mice

All methods were performed in accordance with the relevant guidelines and regulations and animal experiments were approved by the Institutional Animal Care and Use Committee at Rhode Island Hospital (RIH), Providence RI under protocol #5044-21. Animals were three male TRPV1-ChR2-EYFP mice, aged 2-24 months, bred at RIH using the TRPV1-Cre and Ai32(RCL-ChR2(H134R)/EYFP) commercially available lines from Jackson Laboratories. Mice were housed in groups of two or more, unless they received chronic tetrode implants, in which case they were housed in single cages. All experiments were performed using male mice to limit the sex-related variables. Future studies will incorporate female mice to investigate sex-differences in cortical dynamics of somatosensation.

### 2.2 Chronic extracellular electrophysiology

Microdrive implants consisted of between 4-16 tetrodes using either the Neuralynx Halo-5 drive or the Open Ephys shuttleDrive [37]. Tetrodes were made by twisting pairs looped pairs 12.7-micron Nickel-Chrome wire with a polyimide coating (Sandvik) with an open-source tetrode twister (Open Ephys). Depending on initial impedance, tetrodes were electroplated with gold solution (Neuralynx) using a Nano-Z (White Matter LLC) to ensure impedances were between 150-350kOhm; plating was performed with 2 runs at −0.05uA, 1004Hz, with a 2s interval and 1s pause to a 250kOhm target. Drive bodies were coated in a conductive paint (MG Chemicals, Super Shield), which were tied to the ground position on the electrode interface board, for electrical shielding.

All tetrodes were lowered initially between 600-900 microns to account for the thickness of the skull and ensure contact with the cortex. Then, tetrodes were lowered to achieve roughly an equal number of tetrodes in three target zones within S1HL; superficial layers (L2/3), middle layer (L4), and deep layers (L5/6). Approximate estimates for these depths were determined using stereotaxic and histological coordinates. Tetrodes were lowered roughly 20-50 microns/day so that damage to the cortex and surrounding neurons was minimized.

Tetrode recordings were sampled at 30kHz using the Open Ephys acquisition board and low-profile SPI headstage (https://open-ephys.org/). Neural data used for time frequency analysis was filtered between 0-150Hz, and down sampled to 2kHz.

### 2.3 Stereotaxic surgeries

Mice were anesthetized using Isoflurane. Their heads were shaved and sterilized using betadine and 70% ethanol. Mice were then fixed in a stereotactic frame (Kopf instruments). An incision was made down the midline of the scalp and the skin was retracted. The exposed skull was then cleaned and scored to provide a dry surface for bonding with the dental cement. The craniotomy site for implantation was identified by using stereotactic coordinates for S1HL (−0.5 caudal, 1.5 lateral with respect to bregma). A roughly ∼2-4mm craniotomy and 2-3 pilot holes were drilled using a micro drill (Stoelting). Stainless steel screws (000-120, Antrin miniature) were screwed into the pilot holes to provide structural support of the tetrode microdrive as well as provide a ground connection. Following the craniotomy, a durotomy was performed to allow tetrodes to enter the cortex without being damaged and covered with saline for the remainder of the procedure. Custom titanium or stainless-steel head posts were then properly aligned and fixed to the skull using dental cement (MetaBond). The tetrode Microdrive was then slowly lowered over the craniotomy until the base of the drive was flush with the surface of the skull. Dental cement was applied again to secure the drive to the skull. Mice were then placed under a heating lamp or on a warming pad until they were bright, alert, and responsive. Behavioral testing and water restriction began no less than one week following implantation.

### 2.4 Behavioral testing

Prior to behavioral testing, mice underwent at least one thresholding session which consisted of 80 trials of 5 different optogenetic stimulus intensities (10ms pulse width, 0.39-1.18mW/mm^2^), for 400 trials total. Threshold optogenetic stimulus intensity was then determined by the stimulus intensity that evoked a roughly 50% hit rate as determined by scoring hind quarter movements. All behavioral testing sessions consisted of 500 trials that were randomly distributed to ensure that the trial order did not affect the animal’s behavioral response. Mice received four different stimulus types: 100 trials of suprathreshold optogenetic stimulation (10ms pulse width, 4mW/mm^2^), which was determined from previous psychometric responses to optogenetic stimulation [3], 200 trials of threshold optogenetic stimulation as determined by the aforementioned methods (10ms pulse width), 100 trials of tactile stimulation (one cycle of ∼120Hz sine wave), and 100 trials of simultaneous threshold optogenetic and tactile stimulation. Each trial had randomly distributed ISIs (between 3-6s).

### 2.5 Time-frequency analysis

Time-frequency analysis (TFR) was performed by convolving time-series data with a 7-cycle Morlet wavelet over 1-150Hz. To compare power across tetrodes the percent change from baseline was calculated for each trial in every session across all tetrodes within frequency bands of interest by using the 500ms pre-stimulus window.

### 2.6 Spectral Event Analysis

Spectral event analysis was performed using the SpectralEvents toolbox (https://github.com/jonescompneurolab/SpectralEvents) developed in the lab of Dr. Stephanie Jones. Spectral events were identified using the main method implemented in Shin et. Al., 2017; by thresholding peaks of local maxima in un-normalized spectrograms in frequency bands of interest [24]. Spectrograms were calculated using the same method applied in the TFR analysis (see above methods).

### 2.7 Custom force plate

To track hind paw forces during behavioral testing, a load cell was connected to a custom floating footplate onto which the hind paw was restricted. The load cell was connected to a custom circuit that amplified the output voltage of the load cell to be synchronized with the neural data.

### 2.8 Force data collection and analysis

Force data was collected using a custom force plate connected to a 100g load cell (Sparkfun, TAL221) that was integrated into the behavioral platform. The load cell is made up of 4 strain gauges organized into a Wheatstone bridge, which measures force by comparing changes in resistance between the strain gauges. Output from the load cell was amplified by an instrumentation amplifier (Texas Instruments, INA125P) incorporated in a custom circuit board. Output from the load cell amplification board was sampled at 30kHz through the analog input of the Open Ephys acquisition board. Data was filtered between 1-100Hz, and down sampled to 2kHz for analysis. For the purposes of display, force data was filtered between 1-20Hz.

### 2.9 Force subspace generation

Within each session, force responses were z-scored across all trials. As a stimulus reaction could be identified from either a positive or negative deflection in force read out from the load cell, we identified reaction times by setting a z-scored force threshold as being ≥ 1 or ≤ −1. We then took the nearest local minimum preceding the first threshold crossing as the reaction time for that trial.

For generating the force clusters, force data was first visually inspected to identify trials of high pre-stimulus movement, which was rejected from analysis. Force data was then normalized with respect to the 500ms pre-stimulus baseline. As the tactile stimulus generated a large transient in the force response (due to the reducing downward pressure on the force plate by contacting the hind paw), features were extracted from the baselined data 10ms after stimulus onset to ensure that the unsupervised feature extraction and clustering did not mistake the stimulus artifact as part of the reflexive response. Importantly, no behavioral responses occurred within this 10ms window, so excluding this time window from analysis did not bias the reaction time or force subspace analysis. T-distributed stochastic neighbor embedding (tSNE) was then used on the feature subspace to generate clustered data. The first step of tSNE is to generate a high-dimensional map by creating a gaussian distribution for each data point that describes its proximity to all other data points in the set. The second step is to create a low-dimensional map by randomly ordering the high dimensionality data into a lower dimensional space, and then creating a student t-distribution on the low-dimensional data points. Finally, to obtain the low-dimensional space describing the data, the Barnes-Hut algorithm is used to minimize the distance between the high dimensional and low dimensional mapping. K-means clustering is then performed on the low-dimensional data set with a cluster size of k=3. K-means clustering aims to minimize the overall within cluster variation.

### 2.10 Gamma event force distributions

To generate the force distributions in **Figure 5** we first z-scored all force data for each session in every animal. Z-scored force values were then taken at 0, −20, and +20ms with respect event times on corresponding trials, creating a single distribution for each tetrode for every session. Each distribution was centered by subtracting each data point by the mean of the original distribution to get a mean of 0. Bootstrapping was used on the centered distributions to create 10,000 new data sets that were used to create a distribution of z-scored force means.

### 2.11 Statistics

Data was tested for normality using the Anderson-Darling test. If the null hypothesis was rejected (*p*<0.05), then nonparametric statistical tests were used; the Wilcoxon signed-rank test was used for all paired data, the Mann-Whitney U was applied to all unpaired data, and the Kruskal-Wallis test was used to identify significance across all stimulus conditions (suprathreshold, tactile, and strong and weak-threshold). A bootstrap test was used to determine if there was a significant correlation between hind paw force and gamma events for each tetrode on every recording session (see methods on **Gamma event force distributions**).

## 3. Results

### 3.1 Temporally precise surrogate of reflexive response

To achieve temporally precise measurements of behavioral responses, we modified our previous behavioral paradigm [3] to include a custom force plate that enables the measurement of hind paw forces following sensory stimulation as a continuous analog voltage sampled at 30kHz (**Fig. 1A and Supplementary Fig. S1**). This ensured that our behavioral read out could not only keep up with the millisecond timescales inherent for stimulus delivery and neural recording but could also be used to compare with previous experimental results investigating gamma band oscillations that use reflexive measures as the behavioral output. By delivering stimuli through an aperture in the force plate, we tracked the force response to both noxious and innocuous stimuli throughout the course of behavioral testing (**Figs. 1B and C**). The automation and reproducibility of the behavioral paradigm enabled animals to be tested over the course of 500 trials with four distinct stimulus conditions. Suprathreshold optogenetic stimulation and tactile stimulation were used to investigate distinct states of pain and touch, respectively; threshold optogenetic stimulation was used to examine pain saliency, and simultaneous threshold optogenetic and tactile stimulation were used to examine the effect of multi-modal integration.

**Figure 1.**
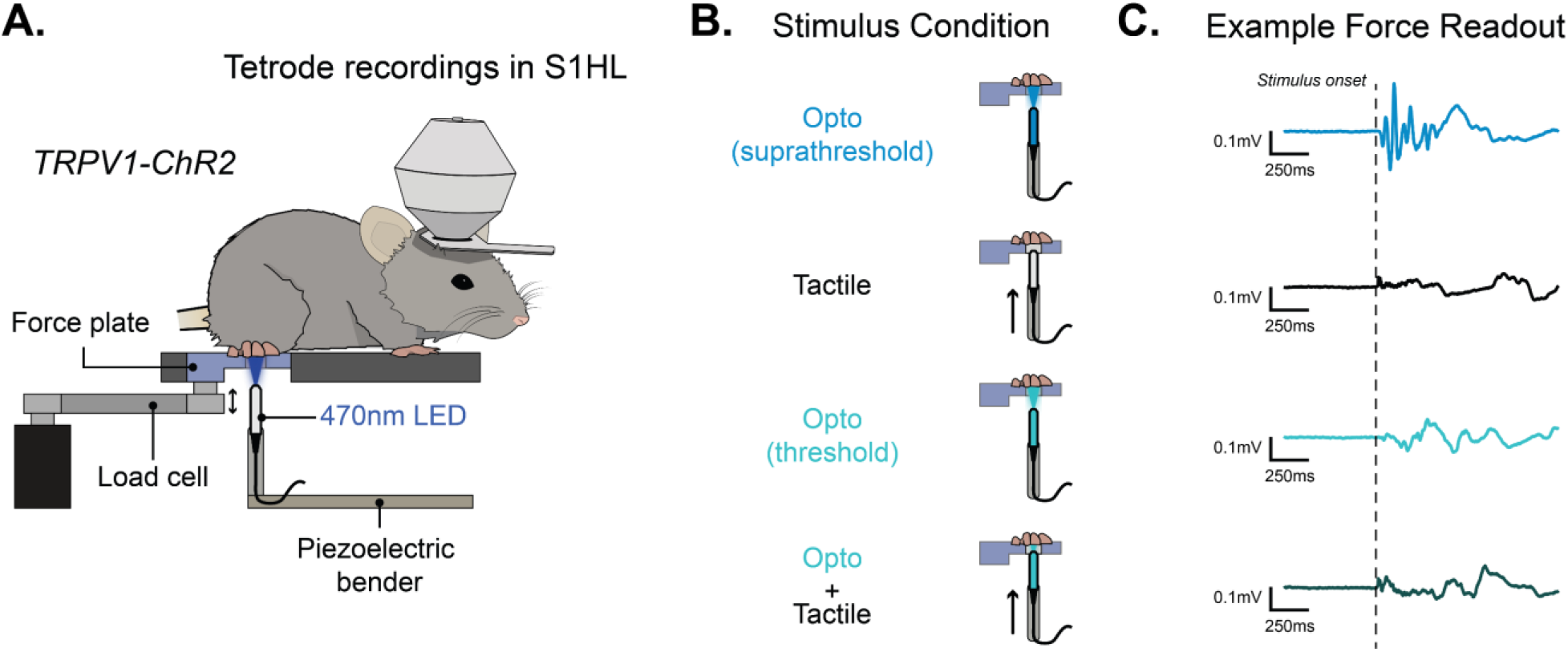
High-speed monitoring of reflexive force response in transgenic mice. (**A**) TRPV1-ChR2 mice implanted with tetrode microdrives were head fixed, and had their hind paw restrained over the force plate through which tactile and optogenetic (470nm LED) stimulation were delivered. (**B**) Mice received 4 stimulus conditions: suprathreshold optogenetic stimulation, tactile stimulation, threshold optogenetic stimulation, and simultaneous threshold optogenetic and tactile stimulation. (**C**) Corresponding representative single trial force responses for each stimulus condition.

### 3.2 Unsupervised hind paw force clustering delineates sensory responses

We then sought to categorize the force responses for each trial within each session in an automated and observer-independent manner. This was achieved by first identifying features in the force read out that correlated with hind paw nocifensive responses and that maximized the difference between tactile and suprathreshold optogenetic trials. These features consisted of percent change in power spectral density (PSD) between 1-30Hz to correlate with paw shaking, maximum and minimum amplitude in the initial phase of the force response (0-65ms) and maximum and minimum amplitude in the later phase of the force response (65-1000ms) to correlate with paw withdrawal height, and maximum rate of change to correlate with paw velocity (**Fig. 2B, *left***). After these features were extracted from each trial within a session, we performed dimensionality reduction using t-distributed stochastic neighbor embedding (tSNE) to reduce the force features of each trial from a 6-dimensional space to a 3-dimensional space (**Fig. 2B, *middle***). We then used K-means clustering with a cluster size of 3 to represent the behavioral response range, which we labeled as ‘strong’, ‘moderate’, and ‘weak’ force subspaces based on their feature space (**Fig. 2B, *right***) and assign scores to each trial (**Fig. 2C**). The *strong force subspace* contained the highest density of suprathreshold optogenetic trials, the *weak force subspace* contained the highest density of tactile trials, and the moderate force subspace contained a mix of threshold and suprathreshold optogenetic, and tactile trials. For all our subsequent analyses, we compared trials in the *strong force subspace* with trials in the *weak force subspace* as this indicated the largest deviation in behavioral response, and a clear distinction between pain and tactile states. Comparisons were made between the *strong* suprathreshold optogenetic trials (supra_str_) and the *weak* tactile trials (tactile_wk_) to characterize differences in sensory modality, and comparisons made between *strong* threshold optogenetic trials (threshold_str_) and *weak* threshold optogenetic trials (threshold_wk_) were used to characterize differences in evoked and non-evoked threshold behaviors (**Fig. 2D**). Moreover, comparisons made between *strong* simultaneous trials (SMT_str_) and *weak* simultaneous trials (SMT_wk_) were used to characterize the effect of somatosensory integration on the intensity of behavioral motor responses. Force response latencies were 34.5±8.2ms for supra_str_ trials, 78.8±48.6ms for tactile_wk_ trials, 84.5±26.8ms for threshold_str_ trials, 154.4±56.3ms for threshold_wk_ trials, and 50.4±62.9ms for the SMT_str_ trials, and 60.78±108.4ms for SMT_wk_ trials. The force onset latencies of TRPV1-ChR2 mice to both supra_str_ and threshold optogenetic stimulation conditions were within the bounds of hind paw reflex latencies of TRPV1-ChR2 mice as determined by high-speed video [1,4]. Furthermore, there was a significant difference between the onset latency of suprathreshold_str_ and tactile_wk_ conditions, and threshold_str_ and threshold_wk_ conditions (*p*>0.05, Mann-Whitney U test, **Supplementary Fig. S2**), but there was no significant difference in onset latency between SMT_str_ and SMT_wk_ conditions.

**Figure 2.**
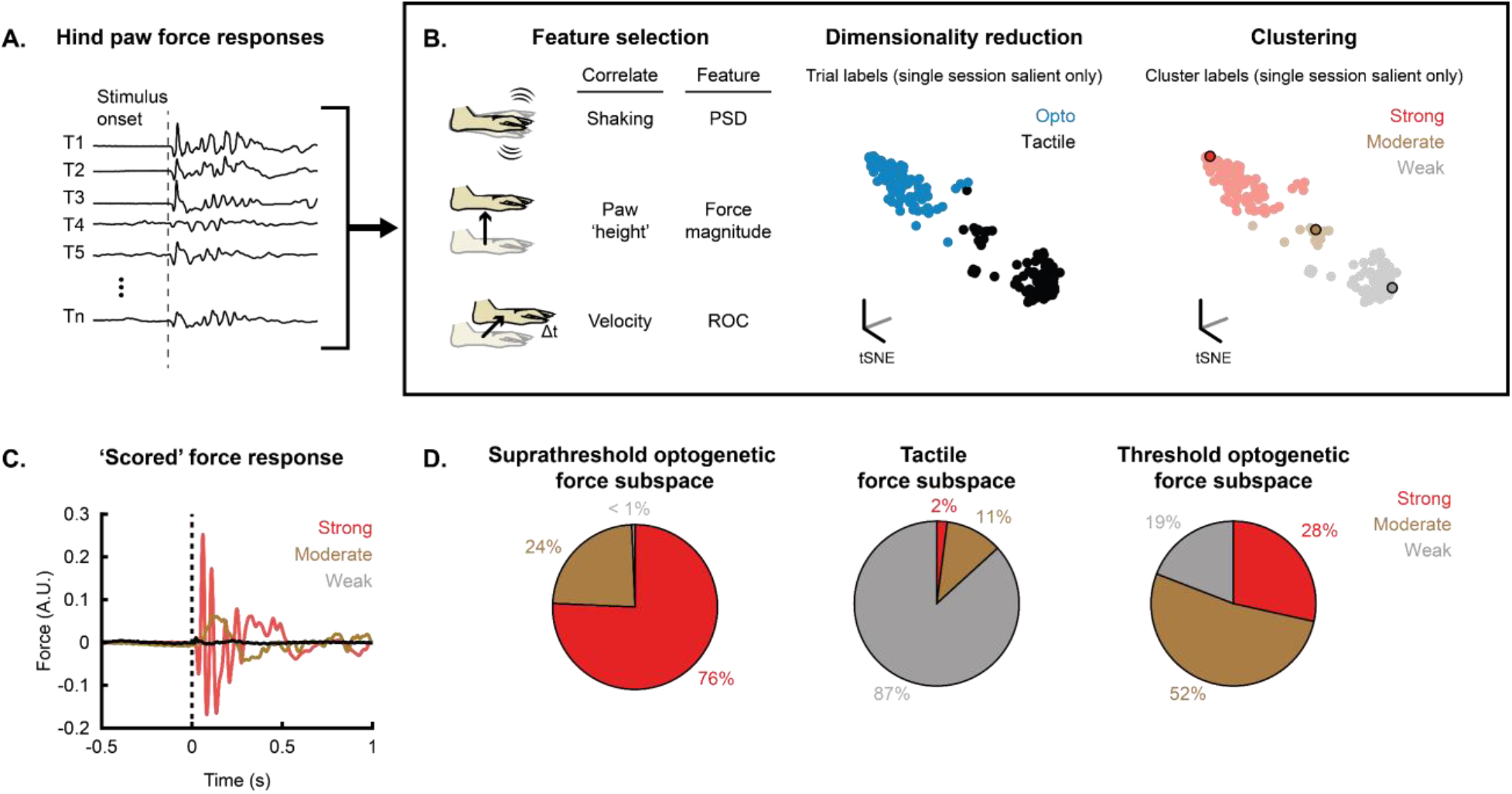
Unbiased force response clustering. (**A**) Example hind paw force responses for different trials. (**B**) Left example illustration of feature selection parameters and corresponding metrics for use in tSNE dimensionality reduction; middle example distribution of trials of a single session (only showing tactile and suprathreshold optogenetic stimulation conditions for simplicity) following dimensionality reduction with tSNE; right k-means clustering of the same data set using a cluster size of k=3. Clusters were identified as strong, moderate, and weak depending on the percentage of suprathreshold optogenetic and tactile trials contained within each cluster. (**C**) Example traces of strong, moderate, and weak force responses taken from **B**. (**D**) Distribution of suprathreshold, tactile, and threshold trials in strong, moderate, and weak force subspaces.

### 3.3 S1 Gamma power increases following peripheral somatosensory stimulation

S1 localized gamma band activity is strongly associated with evoked pain in humans [8,41] and animals [31,40], therefore we sought to determine if gamma band activity was differentially modulated between stimulus conditions. Typically, gamma band activity is defined as a broad range between ∼30-100Hz; however, aspects of cortical processing might be represented by specific sub-bands within this range [26,35]. To account for band-specific activity, we evaluated our neural data in two sub-bands of gamma: low (30-50Hz) and high (70-100Hz). We calculated the time-frequency response (TFR) of the LFP epoched 0.5s prior to, and 1s following stimulus delivery. TFR results revealed that following both tactile, suprathreshold, and threshold optogenetic stimulation, power in high (70-100Hz) and low (30-50Hz) gamma frequency bands increased with respect to the pre-stimulus baseline period (**Figs. 3A-D**).

**Figure 3.**
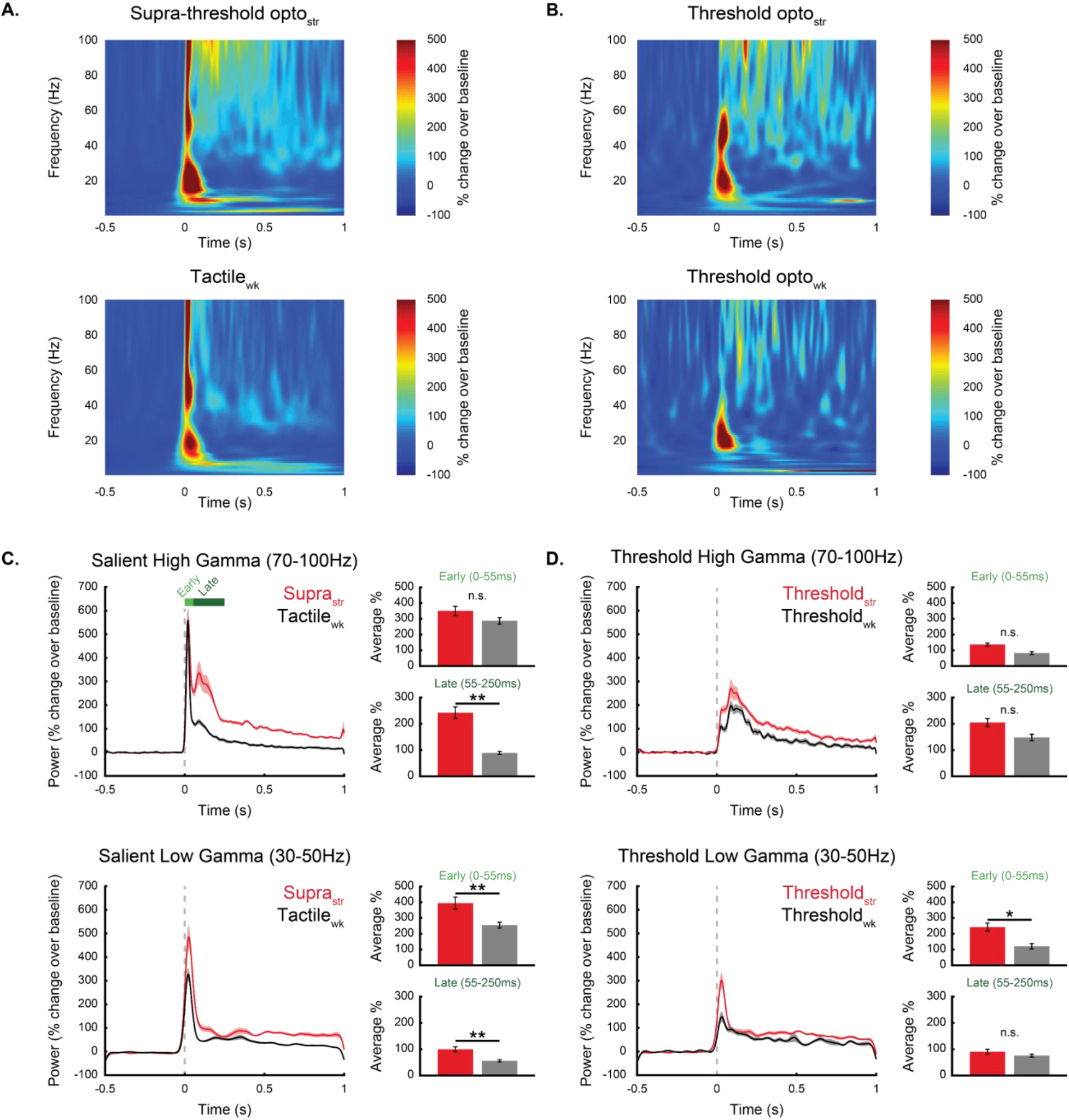
Gamma band activity differs across conditions. (**A-B**) Example average spectrograms from single sessions of force sorted salient and threshold conditions, shown as percentage change from baseline period (−500-0ms pre-stimulus). (**C-D**) Comparisons across all animals between salient and threshold conditions, sorted into high (70-100Hz) and low (30-50Hz) gamma bands. N = 126 samples (measurement from a tetrode on a particular session), **p<0.001, *p<0.05, Kruskal-Wallis test with Tukey-Kramer test for multiple comparisons.

Interestingly, we found that supra_str_ and tactile_wk_ trials exhibited a bimodal response in the high-gamma band, with an early peak occurring between 0-55ms (22.4±3.5ms for supra_str_ and 19.4±2.3ms for tactile_wk_) that was not significantly different between the two conditions (355±201% for supra_str_ and 303±195% for tactile_wk_), and a late peak occurring between 55-250ms (94.7±30.4ms for supra_str_ and 89.6±27.6ms for tactile_wk_) that was significantly greater (*p*<0.001) for the supra_str_ optogenetic condition (232±121%) than for the tactile_wk_ condition (83±32%) (**Fig. 3C, *top***). In the threshold conditions, the bimodal response in the high gamma band was noticeably absent. Although high gamma power increased in the first 55ms following stimulation (threshold_str_: 129±58%, threshold_wk_: 74±59%), it was significantly weaker (*p*<0.001) than the early response for both tactile_wk_ and supra_str_ trials. Only a stronger dominant late peak occurring for both threshold_str_ (201±96%, 103.2±42.5ms) and threshold_wk_ (144±68%, 115.3±38.5ms) trials was present (**Fig. 3D, *top***). Interestingly, there was no significant difference between threshold trials in either the early or late response period; however, both threshold_str_ (*p*<0.001) and threshold_wk_ (*p*<0.05) trials had significantly greater power in the late response when compared to tactile_wk_ trials. This suggests that the early high gamma response may represent absolute stimulus intensity, while the late response may represent modality-specific processing. Both high (70-100Hz) and low (30-50Hz) gamma power increased following stimulation to SMT_str_ and SMT_wk_ trials (**Supplementary Fig. S3**). The response in the high gamma band showed a similar bimodal response as supra_str_ trials, exhibiting an early (0-55ms) and late (55-250ms) component. As the SMT condition consists of both tactile and threshold optogenetic stimuli, it is likely that the early response is representative of the tactile component of the SMT stimulus, while the late response is representative of the threshold optogenetic component of the SMT stimulus. Similar to the supra_str_ and tactile_wk_ trials, there was no significant difference between the magnitude of the early response, but the SMT_str_ late response was significantly greater than the SMT_wk_ late response (**Supplementary Fig. S3B**), further suggesting that the late response is reflective of nociceptive processing.

In the low gamma band (30-50Hz), there was also an early response occurring from 0-55ms that was more pronounced and stereotyped for tactile_wk_ (22.9±6.8ms) and supra_str_ (27.2±7.6ms) trials than for threshold_str_ (59.2±81.6ms) and threshold_wk_ (91.8±109.5ms) trials. Low gamma power during this early response was significantly greater (*p*<0.001) for supra_str_ trials than for tactile_wk_ trials. However, we found no difference between the magnitude of the early response between the tactile_wk_ and the threshold_str_ optogenetic conditions, but a significant difference between the tactile_wk_ and the threshold_wk_ optogenetic conditions (*p*<0.001). Low gamma power was also significantly greater for the threshold_str_ trials than for the threshold_wk_ trials (*p*<0.05) during the early response period. While there was no distinctive secondary response in the low gamma band, low gamma power between 55-250ms was significantly greater (*p*<0.001) for supra_str_ trials than for tactile_wk_ trials, but there was no difference in the later response between threshold_str_ and threshold_wk_ trials. Unlike the high gamma band, the change in power in the low gamma band for the SMT conditions was not consistent with tactile, suprathreshold optogenetic and threshold optogenetic conditions. Both SMT_str_ and SMT_wk_ trials showed a sharp, early response that was not significantly different between the two conditions. The low magnitude late response was, however, significantly greater (*p*<0.05) in SMT_str_ trials than in SMT_wk_ trials (**Supplementary Fig. S3C**).

### 3.4 Transient gamma events correlate with stimulus modality and nocifensive response

While our initial analyses treated gamma band activity as continuous, we noticed that on individual trials gamma band activity in the TFR appeared as transient ‘spectral events’. This raised the question of whether these transient events were responsible for the differences in power in the gamma band shown between trials in the *strong* and *weak* force subspaces. To test this, we implemented a toolbox that identifies transient spectral events in a frequency range of interest from the TFR of time-series data using one of three different algorithms (https://github.com/jonescompneurolab/SpectralEvents). We chose this toolbox specifically as it computes features of interest that help account for increased power observed during trial averaging, like the average TFR shown in **Figure 1**. For our analysis, we implemented an algorithm (method 1) used previously [24] that identifies spectral events as suprathreshold regions within local maxima of the un-normalized TFR. We compared the average event count, onset timing, power, duration, and frequency span of high and low gamma events between supra_str_ and tactile_wk_ trials, as well as threshold_str_ and threshold_wk_ trials for all tetrode recordings (**Fig. 4**). In the high gamma band, we found significant differences between conditions for average event count, onset timing, and duration (*p*<0.05), but not for average event frequency span or maximum power. Of the three significant features, the average event count had the most striking difference when comparing supra_str_ (4.3±0.59 average events/trial) and tactile_wk_ (1.9±0.56 average events/trial) trials, with only one tetrode consistently showing a similar event count between conditions (**Fig. 4A, *top***). This trend also appeared when comparing threshold_str_ (3.8±0.56 average events/trial) and threshold_wk_ (2.55±0.75 average events/trial) trials (**Fig. 4A, *bottom***), however, to a lesser extent. We found similar trends across gamma event features for SMT conditions; average event count, average event onset, and average event duration were significantly (*p*<0.05) different between SMT_str_ and SMT_wk_ trials, with SMT_str_ trials exhibiting more high gamma events, that occurred earlier, and for longer than high gamma events in SMT_wk_ trials (**Supplementary Fig. S4**). We also compared these features between SMT and the threshold optogenetic condition. We found that for both trials clustered in the *strong* and *weak* force subspaces, the SMT condition had a significantly earlier average event onset time than the complimentary threshold optogenetic condition, whereas there was no significant difference between the other features (**Supplementary Figs. S4B and C**). These results suggest that noxious stimuli evoke more high gamma events, and that these events occur earlier and last longer than high gamma events following innocuous stimulation.

**Figure 4.**
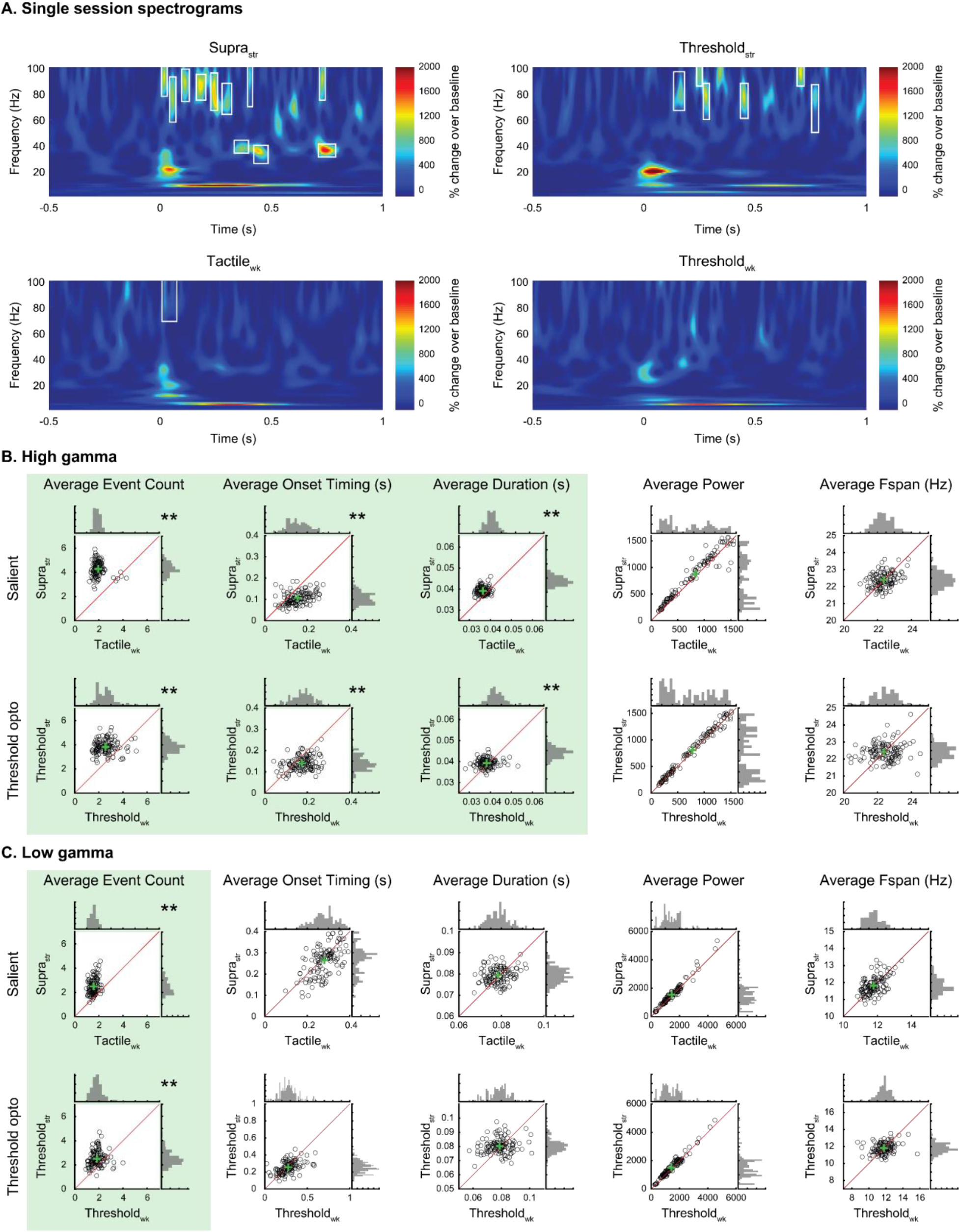
Gamma event features differ between stimulus conditions. (**A**) Single trial examples of spectrograms from different stimulus conditions showing event-like gamma activity (white boxes). (**B**) High gamma (70-100Hz) event comparisons between salient suprathreshold and tactile conditions (top), and strong and weak-threshold conditions (bottom). (**C**) Low gamma (30-50Hz) event comparisons between salient suprathreshold and tactile conditions (top), and strong and weak-threshold conditions (bottom). N = 126 samples (measurement from a tetrode on a particular session), green ‘+’ represent intersection of average event feature for each condition. (n = 126 samples, **p<0.001, two-sided Wilcoxon signed-rank test).

**Figure 5.**
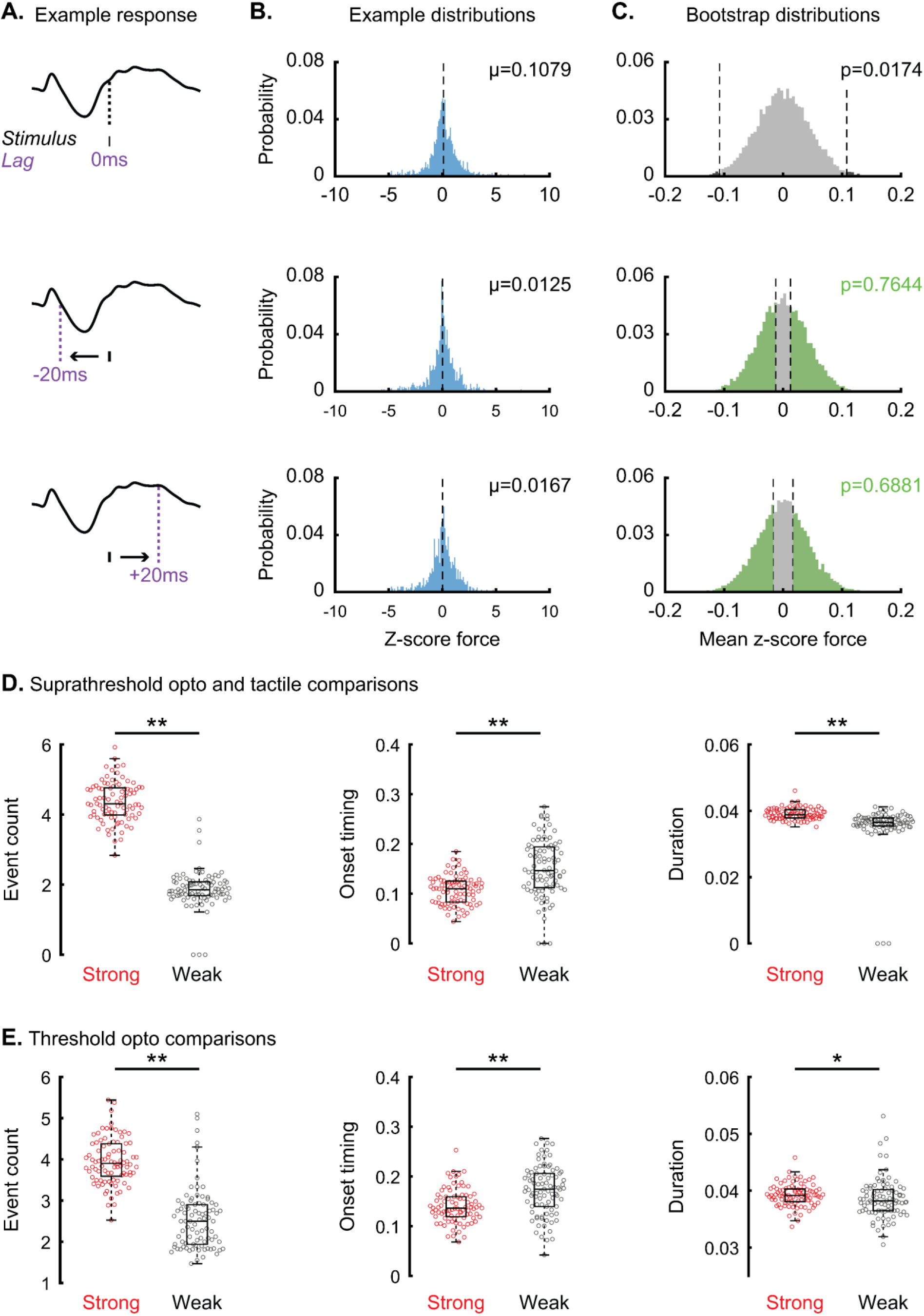
Gamma event difference persist following movement correction. (**A**) Illustration of force responses relative to gamma event used for generating force distributions. (**B**) Example distributions generated from a single tetrode for one session. (**C**) Bootstrapped distributions for the corresponding force distributions in **B**, generated by calculating the mean of 10,000 bootstrapped samples. (**D**) Salient comparisons between strong (suprathreshold optogenetic) and weak (tactile) conditions for previously identified significant features of event count, onset timing, and duration. (**E**) Threshold comparisons between strong and weak conditions for previously identified significant features of event count, onset timing, and duration. (n = 88 samples, **p<0.001, *p<0.05, two-sided Wilcoxon signed-rank test).

In the low gamma band, only the average event count was significantly different (*p*<0.05) between supra_str_ (2.51±0.67 average events/trial) and tactile_wk_ (1.53±0.36 average events/trial) trials as well as threshold_str_ (2.52±0.63 average events/trial) and threshold_wk_ (1.84±0.50 average events/trial) trials. While tactile_wk_ trials appeared to evoke a similar number of low and high gamma events (∼1-2 events/trial on average), supra_str_ trials evoked nearly twice as many events on average across tetrode recordings in the high gamma band than in the low gamma band. This trend also appeared in the comparison between threshold_str_ and threshold_wk_ trials. For the SMT condition, there was only a significant difference (*p*<0.05) in average event count between SMT_str_ and SMT_wk_ trials (**Supplementary Fig. S5A**). Additionally, we found no significant differences for any low gamma event features between SMT and threshold conditions (**Supplementary Figs. S5B and C**).

### 3.5 High Gamma events are not significantly modulated by hind paw movement

One potential confound comparing conditions based on the magnitude of the force read out is that changes in the hind paw force suggests modulation of low-threshold mechanoreceptors throughout the skin that respond to innocuous sensory input such as pressure, touch, and vibration [2]. As we have shown, tactile stimulation elicits increases in gamma band activity in S1, therefore increased movement following noxious stimulation may activate mechanoreceptors, which in turn could elicit gamma events. Although the increase in gamma power in the LFP preceding the onset of the hind paw force response across all conditions suggests that gamma occurs separate from movement, we wanted to determine if hind paw movement was responsible for generating the increase in gamma events. As the average number of events per trial was greater in the high gamma band then in the low gamma band for supra_str_ trials, we analyzed the relationship between the force response and the high gamma events.

We hypothesized that if a change in hind paw force elicited high gamma events, we would expect the distribution of force magnitudes preceding gamma events to have a non-zero mean. To test this, we generated force distributions for each tetrode recording by finding the force magnitude 20ms prior to the timing of the maxima of every high gamma event across all trials (**Fig. 5A**). 20ms was used as this was roughly the delay to maximal high gamma power following peripheral tactile stimulation. We then performed a bootstrapping procedure to generate distributions of average force magnitudes (*see Materials and Methods*). Interestingly, we found that most tetrode recordings (70%) did not show a significant difference between the force distributions and the bootstrapped distributions (**Supplementary Table 1**), which suggests that force magnitude was not responsible for driving spectral events in the high gamma band. We also performed this analysis using time lags of 0 and +20ms, to determine if the force response was consistent around or after gamma events. Like the −20ms lag comparison, most of tetrodes across recording sessions showed no significant difference in force magnitude at 0 (61%) and +20ms (52%) lags. Using this information, we re-computed differences in the high gamma event features, leaving out the tetrode recordings that showed significant differences in the −20ms lagged force distributions (**Figs. 5A** and **B**). After removing these samples, we found that the significant difference in event count between both supra_str_ and tactile_wk_ trials, as well as threshold_str_ and threshold_wk_ trials persisted (**Figs. 5D and E**). In the SMT conditions, we found that the average event count and average event duration maintained a significant difference between SMT_str_ and SMT_wk_ trials, while the significant difference in average event onset time disappeared following the bootstrapping correction (**Supplementary Fig. S6**). Comparisons between SMT and threshold conditions maintained their pre-corrected trend, where the average event onset timing was significantly earlier for the SMT conditions than for the threshold conditions. However, the significance of the average event onset timing in the *strong* force subspace decreased (from *p*<0.001 to *p*<0.05).

## 4. Discussion

In this study, we build on existing literature that investigates neural dynamics in S1 at the level of extracellular field potentials that correlate with noxious and innocuous states that are distinct. Using a novel behavioral metric predicated on evoked hind paw forces, we show that innocuous tactile and noxious stimulations elicit unique responses in the gamma band. Further investigation into the spectral activity revealed that somatosensory stimulation elicits transient events in the gamma range, and that these events occur at a greater rate following noxious stimulation than tactile stimulation. These results provide a deeper understanding of the relationship between gamma activity and sensory processing, and may provide utility to the clinical examination of pain.

### 4.1 Gamma oscillations and sensory processing

Studies show conflicting evidence about the role of gamma band activity in S1; results across animals and humans suggests that gamma band activity is a neural correlate of pain [7,8,20,30,38–41], while other findings suggest that gamma band activity is a general marker of sensory processing [15,17,18,29]. However, as gamma broadly encompasses 30-100Hz oscillations, it is possible that specific types of activity within the gamma range reflect the processing of different features of somatosensory stimulation. We found that both noxious optogenetic stimulation and tactile stimulation yielded increases in power in both high and low gamma bands, which reflect a mechanism of general sensory processing. However, when analyzing high frequency activity as spectral events, we found that the number, duration, and timing of transient gamma events correlated with suprathreshold and threshold optogenetic stimuli that elicited *strong* force responses (i.e. nocifensive). These results show that certain features of gamma band activity may differentially reflect noxious and innocuous processing, a conclusion that is consistent with results from human EEG showing differential activation within the high gamma band in response to noxious laser and innocuous tactile stimulation [15].

To understand how gamma activity may reflect the processing of different modalities of somatosensation, it is important to understand the circuit mechanisms that drive extracellular gamma oscillations in S1. High frequency oscillations have been shown to occur intrinsically within cortical areas through networks of inhibitory-inhibitory neuron coupling or excitatory-inhibitory neuron coupling [5,21,34], or through gamma oscillatory feedforward drive [9,33]. Recent studies have linked fast-spiking inhibitory interneurons to the generation of gamma oscillations in S1 following noxious stimulation [30,40]. However, as identifying the circuit mechanisms of gamma oscillations was outside the scope of these experiments, future studies would require the implementation of cell-specific manipulations, such as optogenetic activation or high-speed voltage imaging of inhibitory interneuron subpopulations.

Anatomical evidence suggests that feedforward drive may be partially responsible for differences in S1 responses elicited by noxious and innocuous stimulation. The spinothalamic tract (STT), which relays noxious information from the periphery, sends synapses from thalamus to superficial S1, while the dorsal column-medial lemniscal (DCML) pathway, which relays innocuous information from the periphery, sends thalamic projections to middle layers of S1 [36]. The target divergence may mean that noxious and innocuous information may produce different signatures of gamma activity by driving the same S1 circuitry in a laminar-specific fashion. This idea has been recently explored with a biophysically principled neural model of S1 that shows 40Hz gamma drive to the distal dendrites of pyramidal neurons in superficial layers is sufficient to recreate features of noxious laser-induced evoked responses measured in humans [32].

### 4.2 Movement induced gamma

One potential confound to our results is that comparisons between trials are based on the magnitude of the hind paw force response, which might indicate that the paw movement elicits innocuous sensory stimulation that underlies the generation of gamma activity. Although previous findings show that increases in gamma band power induced by noxious stimulation were not a result of stimulus driven movements [30], we needed to determine if this was the case the generation of gamma events. We found that increases in gamma power occurred prior to the initiation of hind paw movement as determined by the force read out. We also found that across sessions, high gamma events on most tetrodes displayed no dependence on preceding hind paw forces, which suggests that hind paw forces experienced during behavioral testing were insufficient to elicit the gamma events investigated in our study.

Another possibility is that gamma events are dependent on complex movement patterns, as cutaneous sensory afferents in the glabrous skin of the hind paw make up a multitude of receptors that respond to skin indentation, stretch, movement, and vibration. Furthermore, these receptors can be rapidly adapting sensory afferents, which respond to the onset and offset of indentation to the skin, and slowly adapting sensory afferents, which respond to continuous indentation of the skin [2]. Although these cutaneous receptors show remarkable organization in both their innervations within the skin and their laminar organization within the spinal cord [2,11], S1 shows less of a discernable somatotopic or laminar distinction among these primary sensory receptors [16]. This problem is further compounded by the fact that we have no way of telling post hoc what sensory receptors were activated following stimulus-evoked movements. Interestingly, active and passive movement have been shown to gate sensory responses in S1 both from bottom-up and top-down mechanisms [22]. Furthermore, passive movement has been shown to gate high-frequency oscillations in S1 at the gamma range [29]. These findings suggest that movement following noxious stimulation initialized by spinal reflexive circuitry and modulated by top-down control could gate sensory drive to S1 that occurred because of movement-induced activation of cutaneous sensory afferents. However, this does not mean that gating of sensory information during movement would entirely prohibit the generation of gamma events in S1, but it may point to a potential mechanism designed to help regulate the processing of specific types of sensory information within the gamma frequency band.

### 4.3 Experimental considerations

It is important to highlight potential differences in experimental setup between our work and prior studies. First, most studies investigating gamma band activity in both mice and humans use noxious thermal (laser) or mechanical (Von Frey) stimulation. Here, we implement time-resolved noxious peripheral optogenetic stimulation. Due to the use of the TRPV1-ChR2 mouse line, the optogenetic stimulus would be most similar to a noxious thermal sensation, as TRPV1 receptors open in response to temperatures in excess of 43°C. However, unlike optogenetic activation, thermal activation of TRPV1 receptors is not instantaneous as it takes time for the skin to reach the temperature necessary to activate channel opening [6,14]. Therefore, the use of optogenetic stimulation might dilate the timeline of the neural response relative to standard thermal stimulation. Additional deviations from standard thermal stimuli might arise because activation of primary nociceptors in TRPV1-ChR2 mice is done through the optically induced opening of ChR2 as opposed to the thermally induced opening of TRPV1. While both ChR2 and TRPV1 are non-specific cation channels and therefore are likely to produce similar intracellular responses, differences in their channel gating could produce dissimilar cellular dynamics that lead to differences in downstream neural activation and resulting behavior. For example, activation of MrgprA3+ containing C-fibers through ionotropic or metabotropic mechanisms yields differing behavioral responses in mice [23]. Therefore, future experiments should directly compare the neural and behavioral evoked responses of optogenetically and thermally mediated activation of TRPV1 containing primary nociceptors in TRPV1-ChR2 mice.

Our results provide further insight into the role of gamma band activity in pain processing by investigating differences in neural activity in S1 evoked by noxious and innocuous stimuli. These results replicate previous findings that shows gamma band activity reflects the processing of both painful and nonpainful information. By further analyzing gamma band activity as spectral events, we were able to identify differences in gamma event features between somatosensory modalities and threshold nociceptive responses. As such, these results better define sensory specific differences in the gamma frequency band that could prove useful in identifying pain in clinical settings.

## 5. Acknowledgements

This work was supported in part through the NIH NINDS BRAIN Initiative under grant 1R01NS108414-01 (C.S. and D.A.B.) and the NIH T32 Training Grant MH115895 (C.B.). The authors declare no conflicts of interest. We would like to thank Dr. Brian Leblanc (Rhode Island Hospital) for administering and maintaining the animal care and use protocol and for his discussions on behavioral testing and electrophysiology, Dr. Jacqueline Hynes and Dr. Hyeyoung Shin for discussions on data analysis, and Dr. Muhammad Edhi for assisting in maintenance of animal colonies.

## Data availability

All data will be freely available upon request to the corresponding author.

