## Supplemental material for "Transient gamma events delineate somatosensory modality in S1"

**Supplemental Figures**

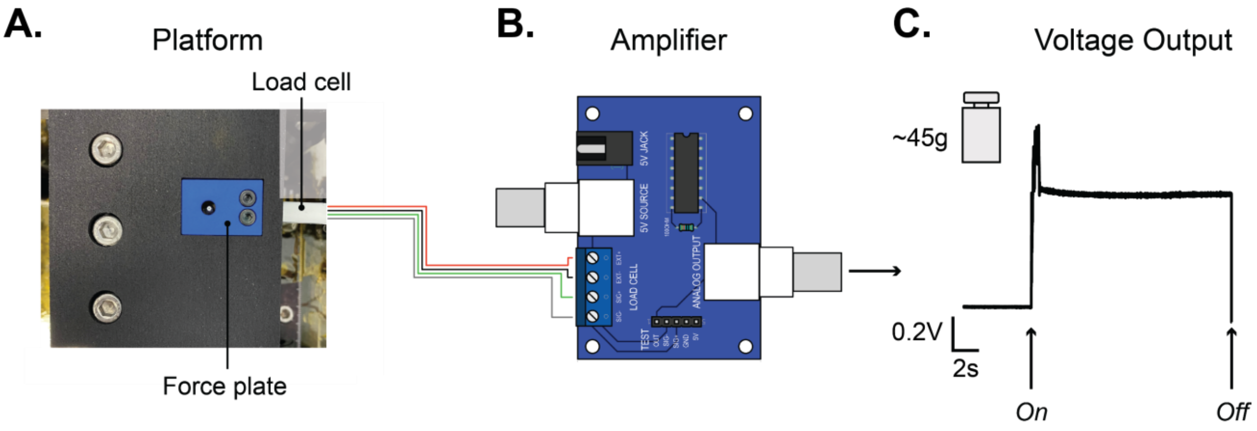


**Figure S1.** Custom force plate design. (**A**) Custom force plate with stimulus aperture connected to load cell. (**B**) Illustration of custom circuit used to amplify analog output from load cell so that it can be registered with the data acquisition system. (**C**) Example voltage response from a ~45g weight, initial spike shows increase in force due to placement and adjustment of weight on force plate, with a rapid settle to baseline output.

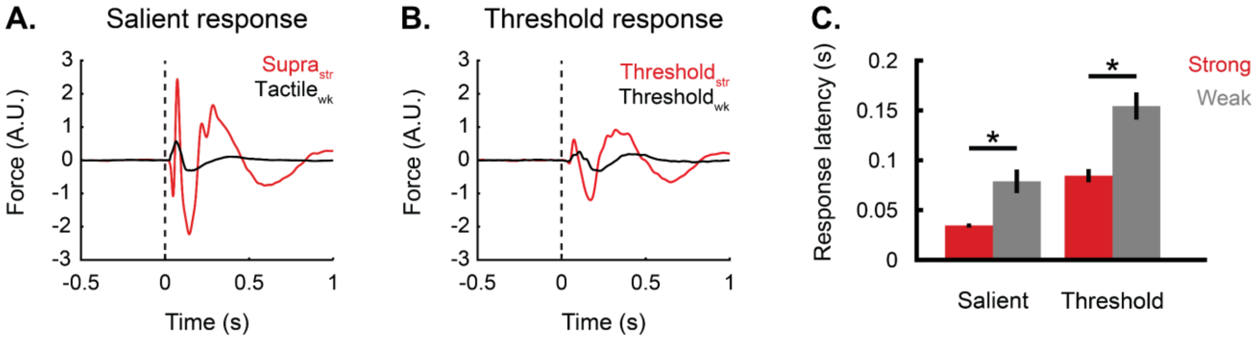


**Figure S2.** Force sorted response comparisons. (**A**) Average z-scored force responses of supra$_{str}$ optogenetic trials (red, n=1287 trials) and tactile$_{wk}$ trials (black, n=1475 trials) across all animals. Representative force responses for supra$_{str}$ (red) and tactile$_{wk}$ (black) conditions. (**B**) Average z-scored force responses of threshold$_{str}$ (red, n=968) and threshold$_{wk}$ (black, n=650) optogenetic trials across all animals. (**C**) Comparison of average force response latency of supra$_{str}$ and tactile$_{wk}$ (salient) conditions and threshold$_{str}$ and threshold$_{wk}$ (threshold) conditions. In each condition, the force response latency for strong sorted trials were significantly faster than the corresponding force response latency for weak sorted trials (*$p<0.05$, two-sided Wilcoxon signed-rank).

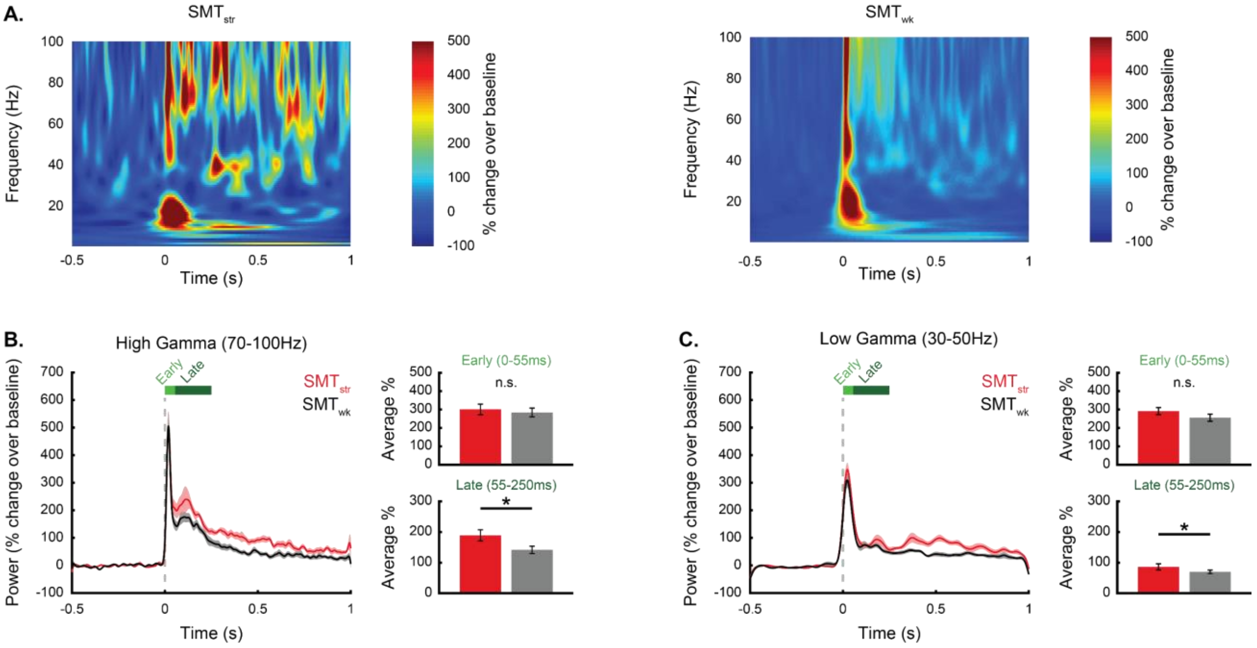


**Figure S3.** TFR and gamma band activity for SMT conditions (**A**) Example average spectrogram showing the percent change in power with respect to the 500ms pre-stimulus baseline for strong (*left*) and weak (*right*) SMT conditions. (**B**) Average percent change over baseline in the high gamma band (70-100Hz) across all tetrode recordings. *Inset* comparison of early (0-55ms) and late (55-250ms) responses. (**C**) Average percent change over baseline in the low gamma band (30-50Hz). *Inset* comparison of early (0-55ms) and late (55-250ms) responses. *$p$<0.05, two-sided Wilcoxon signed-rank test, n=126.

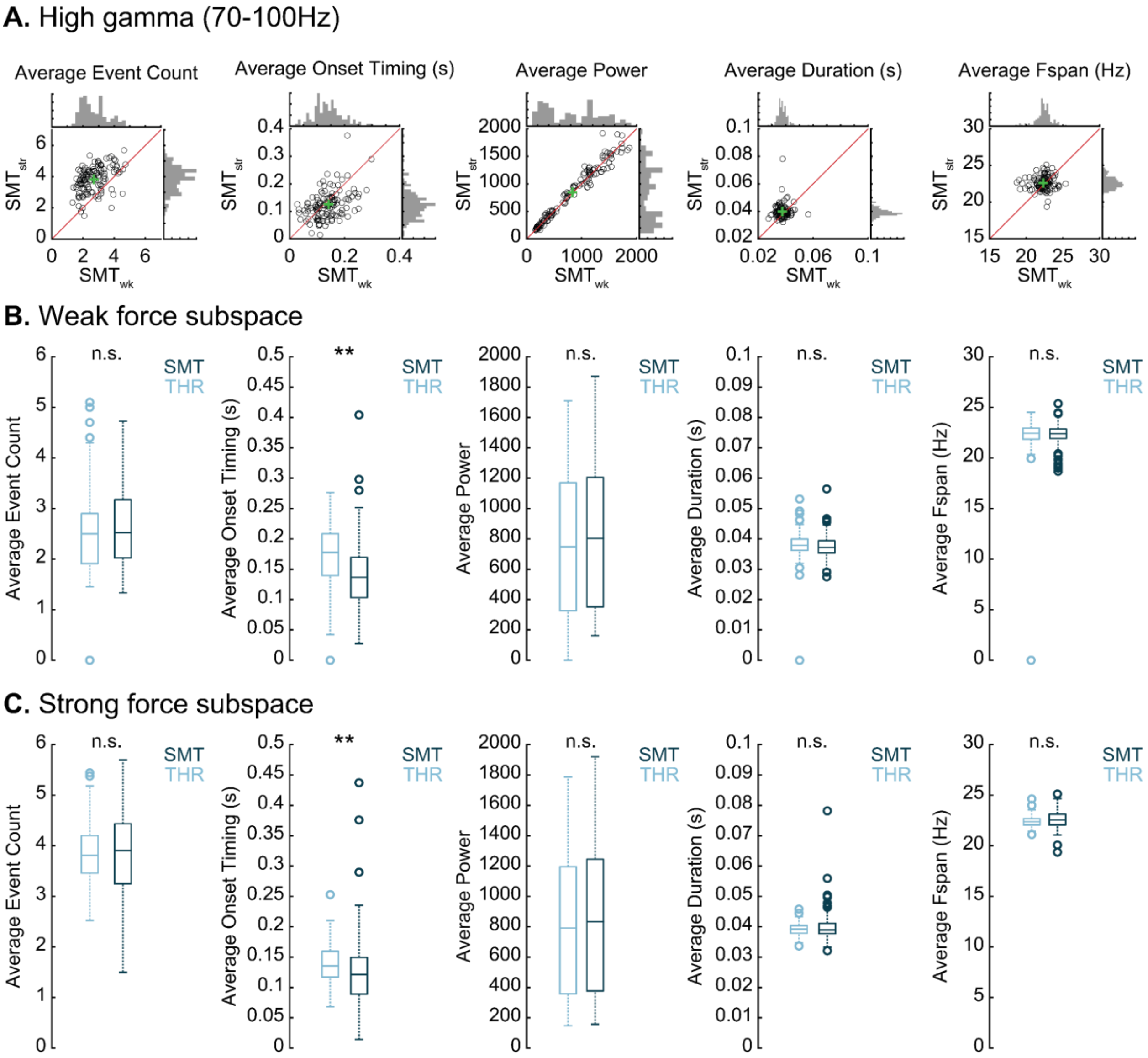


**Figure S4.** High gamma event features. (**A**) Comparison between SMT weak and strong threshold conditions for average event count, average onset timing, average power, average duration, and average frequency span. Green '+' indicates means between both conditions, red line through the origin represents y=x. (**B**) Comparison between SMT and THR conditions in the weak force subspace. (**C**) Comparisons between SMT and THR conditions in the strong force subspace. N.S. $p>0.05$, ** $p<0.001$, two-sided Wilcoxon signed-rank test.

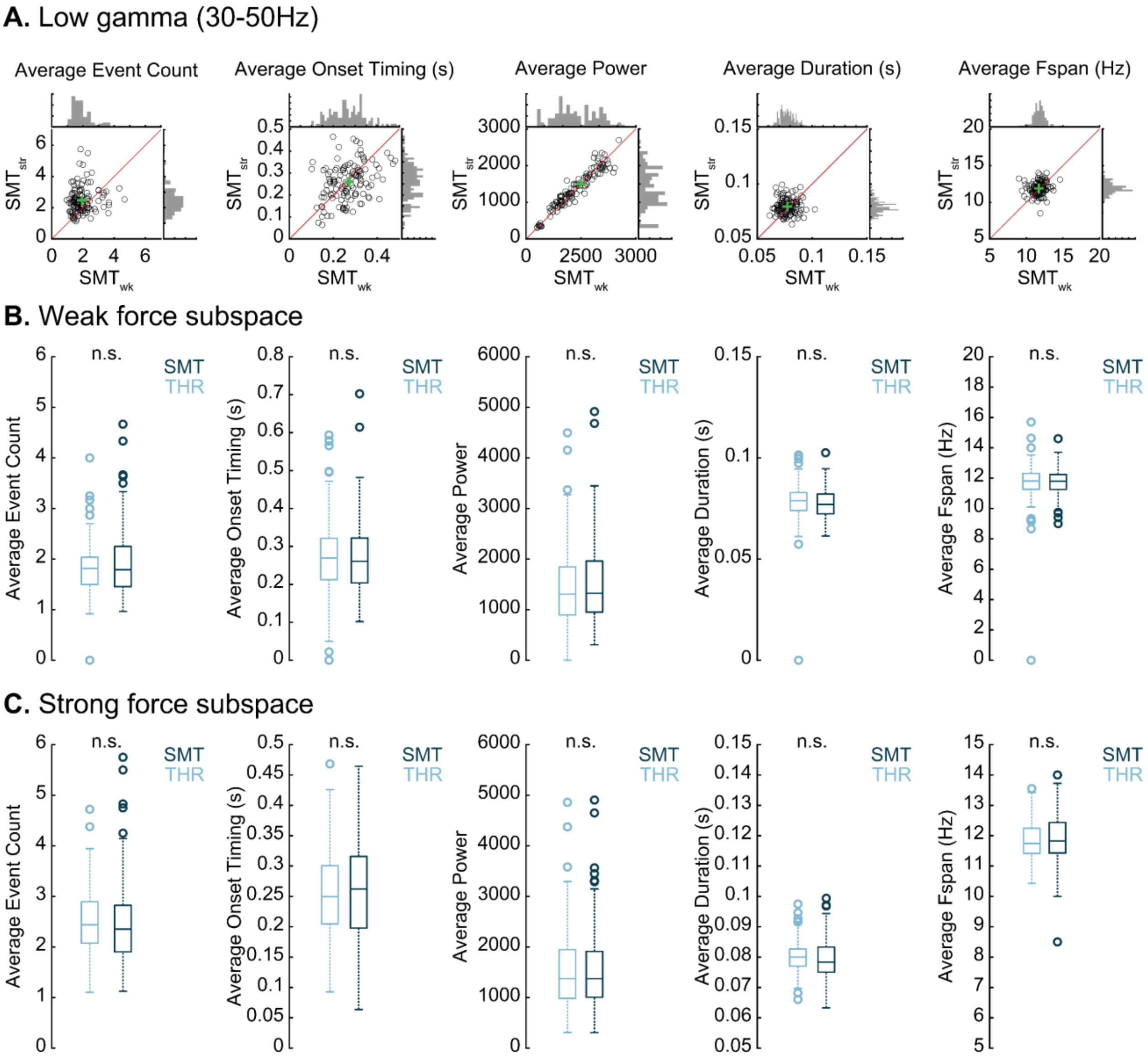


**Figure S5.** Low gamma event features. (**A**) Comparison between SMT weak and strong threshold conditions for average event count, average onset timing, average power, average duration, and average frequency span. Green '+' indicates means between both conditions, red line through the origin represents y=x. (**B**) Comparison between SMT and THR conditions in the weak force subspace. (**C**) Comparisons between SMT and THR conditions in the strong force subspace. N.S. $p>0.05$, two-sided Wilcoxon signed-rank test.

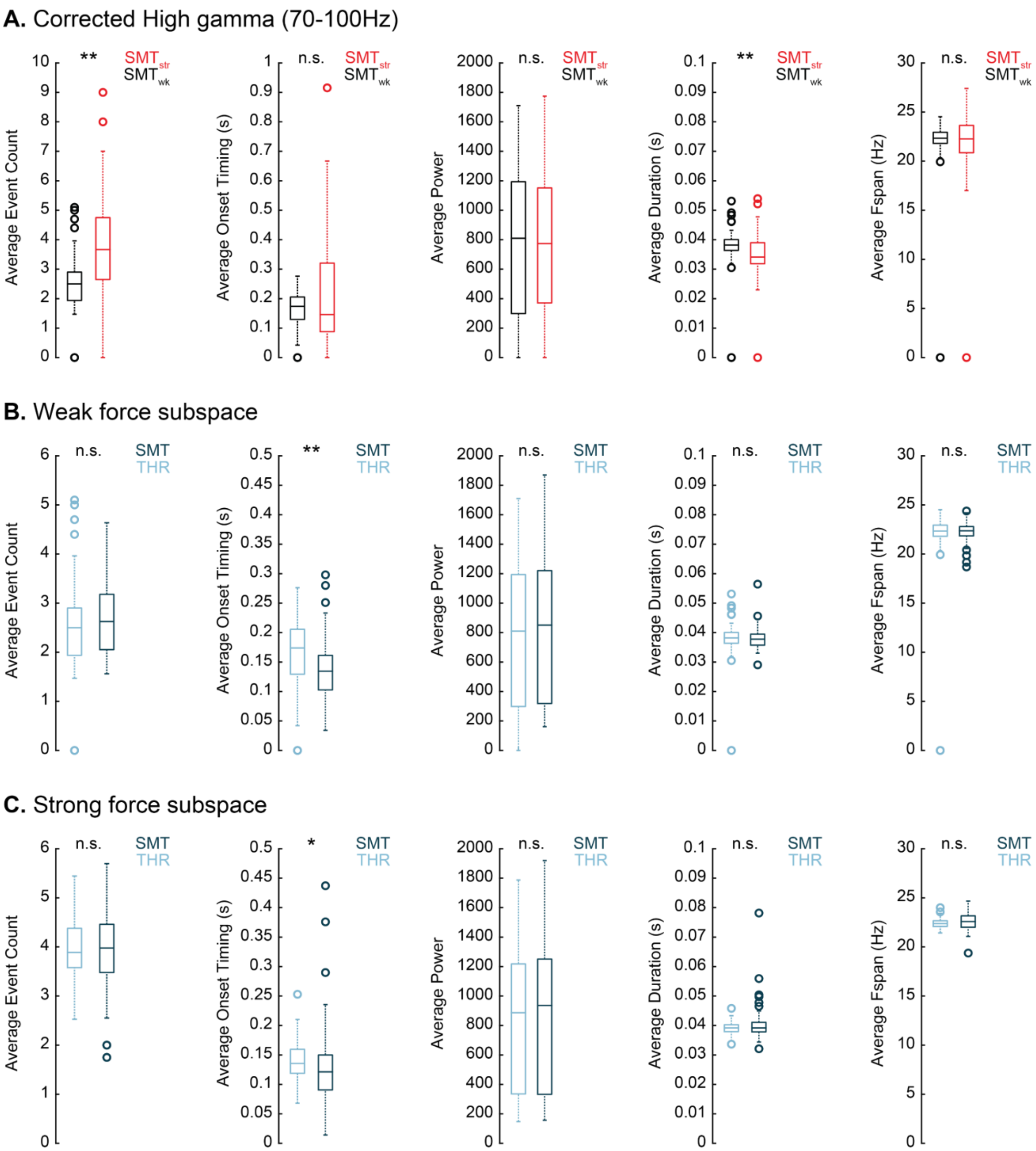


**Figure S6.** Force corrected high gamma event features. (**A**) Comparison between SMT weak and strong threshold conditions for average event count, average onset timing, average power, average duration, and average frequency span. (**B**) Comparison between SMT and THR conditions in the weak force subspace. (**C**) Comparisons between SMT and THR conditions in the strong force subspace. N.S. $p>0.05$, ** $p<0.001$, * $p<0.05$, two-sided Wilcoxon signed-rank test.

### Supplemental tables

### **Table 1.** *p* values for gamma event lagged distributions

#### 0ms lag p-values

| Subject | Tetrode | S1 | S2 | S3 | S4 | S5 | S6 |
|---|---|---|---|---|---|---|---|
| SUBJ1 | T1 | 0.6897 | 0.7194 | 0.3436 | 0.6995 | 0.529 | 0.0790 |
| SUBJ1 | T2 | 0.1447 | 0.9356 | 0.5435 | 0.2579 | 0.0003 | 0.6751 |
| SUBJ1 | T3 | 0.4645 | 0.9945 | 0.7230 | 0.5279 | 0.6301 | 0.1003 |
| SUBJ1 | T4 | 0.1332 | 0.0822 | 0.0037 | 0.2315 | 0 | 0.1651 |
| SUBJ1 | T5 | 0.1294 | 0.0002 | 0.4305 | 0.2768 | 0.0030 | 0.0162 |
| SUBJ1 | T6 | 0.1598 | 0.1426 | 0.6333 | 0.9330 | 0 | 0.4650 |
| SUBJ1 | T7 | 0.2302 | 0.7959 | 0.0162 | 0.0597 | 0 | 0.5540 |
| SUBJ1 | T8 | 0.6163 | 0.1275 | 0.8462 | 0.5890 | 0.4809 | 0.9376 |
| SUBJ2 | T1 | 0.0890 | 0 | 0.2595 | 0.0321 | 0.9426 | 0 |
| SUBJ2 | T2 | 0.8744 | 0.4673 | 0.0127 | 0.4918 | 0.1641 | 0 |
| SUBJ2 | T3 | 0.0035 | 0.0812 | 0.7639 | 0.0006 | 0.3414 | 0 |
| SUBJ2 | T4 | 0.1148 | 0 | 0.9263 | 0.0564 | 0.0780 | 0.003 |
| SUBJ2 | T5 | 0.0218 | 0 | 0.1144 | 0.3756 | 0.0023 | 0 |
| SUBJ2 | T6 | 0.1798 | 0 | 0.9543 | 0 | 0.8246 | 0.0001 |
| SUBJ2 | T7 | 0.0055 | 0.0004 | 0.4369 | 0.1283 | 0.1224 | 0 |
| SUBJ2 | T8 | 0.2982 | 0.9047 | 0.0095 | 0.8888 | 0.8461 | 0.0181 |
| SUBJ3 | T1 | 0.0011 | 0.6356 | 0 | 0.2029 | 0.0983 | |
| SUBJ3 | T2 | 0 | 0.0066 | 0 | 0.0440 | 0.0374 | |
| SUBJ3 | T3 | 0.0085 | 0.0189 | 0 | 0.1218 | 0.9734 | |
| SUBJ3 | T4 | 0.0002 | 0.0069 | 0 | 0.3161 | 0.9469 | |
| SUBJ3 | T5 | 0.0098 | 0.1168 | 0 | 0.1295 | 0.8086 | |
| SUBJ3 | T6 | 0.0106 | 0.2212 | 0 | 0.4233 | 0.0504 | |

#### -20ms lag p-values

Sessions

| Subject | Tetrode | S1 | S2 | S3 | S4 | S5 | S6 |
|---|---|---|---|---|---|---|---|
| SUBJ1 | T1 | 0.5514 | 0.4419 | 0.0194 | 0.1430 | 0.9606 | 0.8482 |
| SUBJ1 | T2 | 0.0176 | 0.9686 | 0.0015 | 0.2601 | 0 | 0.4177 |
| SUBJ1 | T3 | 0.6286 | 0.0750 | 0.0233 | 0.0046 | 0.0217 | 0.5332 |
| SUBJ1 | T4 | 0.0302 | 0.4195 | 0.8916 | 0.9914 | 0.6026 | 0.8037 |
| SUBJ1 | T5 | 0.2356 | 0.1704 | 0.1182 | 0.1543 | 0.5058 | 0.1090 |
| SUBJ1 | T6 | 0.0697 | 0.2926 | 0.9017 | 0.7333 | 0.0355 | 0.5503 |
| SUBJ1 | T7 | 0.8091 | 0.0936 | 0.7590 | 0.2943 | 0.0936 | 0.0658 |
| SUBJ1 | T8 | 0.1859 | 0.1362 | 0.0594 | 0.9618 | 0.0884 | 0.9250 |
| SUBJ2 | T1 | 0.0038 | 0 | 0.1730 | 0 | 0.4686 | 0 |
| SUBJ2 | T2 | 0.9806 | 0.1596 | 0.0203 | 0.0218 | 0.3992 | 0 |
| SUBJ2 | T3 | 0.0334 | 0.0030 | 0.5520 | 0 | 0.1050 | 0 |
| SUBJ2 | T4 | 0.3122 | 0 | 0.5170 | 0.0883 | 0.0789 | 0 |
| SUBJ2 | T5 | 0.4637 | 0 | 0.1369 | 0.0165 | 0.0817 | 0 |
| SUBJ2 | T6 | 0.6849 | 0 | 0.6383 | 0.0001 | 0.2243 | 0 |
| SUBJ2 | T7 | 0.6270 | 0.0004 | 0.4608 | 0.0004 | 0.5428 | 0 |
| SUBJ2 | T8 | 0.7336 | 0.2682 | 0.0003 | 0.2055 | 0.7843 | 0.0338 |
| SUBJ3 | T1 | 0.2241 | 0.1867 | 0 | 0.6165 | 0.0654 | |
| SUBJ3 | T2 | 0.0032 | 0.7166 | 0 | 0.0058 | 0.3213 | |
| SUBJ3 | T3 | 0.0927 | 0.9356 | 0 | 0.0671 | 0.2710 | |
| SUBJ3 | T4 | 0.2523 | 0.4379 | 0 | 0.5887 | 0.4504 | |
| SUBJ3 | T5 | 0.9856 | 0.2375 | 0 | 0.6003 | 0.2598 | |
| SUBJ3 | T6 | 0.5925 | 0.4704 | 0 | 0.4576 | 0.7395 | |

#### +20ms lag p-values

| Subject | Tetrode | S1 | S2 | S3 | S4 | S5 | S6 |
|---|---|---|---|---|---|---|---|
| SUBJ1 | T1 | 0.1562 | 0.0587 | 0.0001 | 0.0043 | 0.0051 | 0.4224 |
| SUBJ1 | T2 | 0.2054 | 0.0161 | 0.0089 | 0.0033 | 0.0178 | 0.7679 |
| SUBJ1 | T3 | 0.0068 | 0.0102 | 0.2412 | 0.9370 | 0.8837 | 0.9042 |
| SUBJ1 | T4 | 0.9534 | 0.1580 | 0.5872 | 0.8253 | 0 | 0.6946 |
| SUBJ1 | T5 | 0.0014 | 0 | 0.0010 | 0.1661 | 0.0023 | 0.6421 |
| SUBJ1 | T6 | 0.5392 | 0.1914 | 0.8147 | 0.3831 | 0.0001 | 0.3156 |
| SUBJ1 | T7 | 0.1749 | 0.1695 | 0.6891 | 0.8500 | 0.0087 | 0.7664 |
| SUBJ1 | T8 | 0.0215 | 0.2575 | 0.0082 | 0.0158 | 0.3111 | 0.3368 |
| SUBJ2 | T1 | 0 | 0 | 0.0093 | 0.0008 | 0.3207 | 0 |
| SUBJ2 | T2 | 0.0467 | 0.1123 | 0.0642 | 0.0600 | 0.8145 | 0 |
| SUBJ2 | T3 | 0.0772 | 0.0035 | 0.6294 | 0 | 0.0003 | 0 |
| SUBJ2 | T4 | 0.3422 | 0 | 0.1769 | 0.0428 | 0.1409 | 0.0006 |
| SUBJ2 | T5 | 0.0006 | 0 | 0.9031 | 0.0148 | 0.4750 | 0 |
| SUBJ2 | T6 | 0.3661 | 0 | 0.5123 | 0.0001 | 0.2933 | 0.0132 |
| SUBJ2 | T7 | 0.0073 | 0.0001 | 0.6163 | 0.0321 | 0.5277 | 0 |
| SUBJ2 | T8 | 0.2034 | 0.9457 | 0.0823 | 0.4667 | 0.0024 | 0.0525 |
| SUBJ3 | T1 | 0.0883 | 0.9813 | 0 | 0.8176 | 0.0982 | |
| SUBJ3 | T2 | 0 | 0.0004 | 0 | 0.5875 | 0.5084 | |
| SUBJ3 | T3 | 0.0082 | 0.0128 | 0 | 0.7397 | 0.7636 | |
| SUBJ3 | T4 | 0.0004 | 0.0193 | 0 | 0.5308 | 0.6382 | |
| SUBJ3 | T5 | 0.0029 | 0.0015 | 0 | 0.4271 | 0.5754 | |
| SUBJ3 | T6 | 0.0084 | 0.1244 | 0 | 0.7076 | 0.8100 | |

removed (*p*<0.05)

remained (p>0.05)
